# Time-Resolved Transcriptomics Reveal Spliceosomal Disruption and Senescence Pathways in Crocin-Treated Hepatocellular Carcinoma Cells

**DOI:** 10.1101/2025.08.05.668798

**Authors:** David Roy Nelson, Amphun Chaiboonchoe, Weiqi Fu, Amnah Salem Alzahmi, Ala’a Al-Hrout, Amr Amin, Kourosh Salehi-Ashtiani

## Abstract

Saffron-derived crocin exhibits anti-cancer properties, but the pathways underlying its effects remain incompletely characterized. Here, we utilized a high-dose perturbation strategy (1–2 mM crocin) to probe maximal pathway engagement in HepG2 hepatocellular carcinoma cells via time-series transcriptomics. We treated cells for 2, 6, 12, and 24 h and analyzed transcriptomic and splicing profiles at each timepoint. We identified 7400–12,100 differentially expressed genes (DEGs) per condition, with the higher dose (CR2) producing more total DEGs but the lower dose (CR1) demonstrating differential pathway prioritization. The spliceosome pathway ranked first among downregulated pathways for CR1 across multiple timepoints (false discovery rate, FDR *p* = 10^−21^ to 10^−36^) but only fourth for CR2, suggesting dose-dependent differences in pathway prioritization. Differential splicing analysis revealed functional spliceosome disruption, with 2000–2600 significant exon skipping events per condition and aberrant splicing of spliceosome components including HNRNPH1 (change in percent spliced in, dPSI = −0.78 to −0.89). Additionally, 66 genes implicated in non-alcoholic fatty liver disease were downregulated at 24 h (FDR *p* = 8×10^−8^). Crocin exposure consistently downregulated spliceosomal machinery genes while upregulating senescence and autophagy pathways. These findings identify spliceosome components and RNA processing machinery as crocin-sensitive pathways.

## 1 Introduction

Saffron (*Crocus sativus* L.) and its bioactive component crocin have demonstrated anti-cancer properties in multiple studies, yet their molecular mechanisms remain incompletely understood [1]. Crocin, a digentiobiose ester of crocetin, suppresses HCC cell survival through various downstream effectors [1]. Studies from our group have demonstrated that saffron extract and crocin exhibit pro-apoptotic, anti-inflammatory, and anti-oxidative effects against HCC, with crocin combined with sorafenib improving tumor inhibition in cirrhotic-HCC models [2].

Hepatocellular carcinoma represents the most common primary liver malignancy and a leading cause of cancer mortality worldwide [3]. Major risk factors include chronic hepatitis B/C infection, alcoholic liver disease, and non-alcoholic steatohepatitis (NASH), with 70–90% of patients having underlying chronic liver disease [4]. Current treatments include surgical resection, transplantation, and transarterial chemoembolization, but novel targeted therapies are urgently needed [5]. Aberrant RNA splicing has emerged as a hallmark of cancer, with splicing dysregulation contributing to oncogenesis in over 90% of malignancies [6, 7]. FDA-approved spliceosome modulators have shown efficacy in hematologic cancers, highlighting the therapeutic potential of targeting this machinery [8, 9, 10].

Non-alcoholic fatty liver disease (NAFLD) represents another critical link between metabolic dysfunction and HCC development. The progression from NAFLD to NASH and ultimately HCC involves coordinated changes in mitochondrial function, lipid metabolism, and inflammatory signaling [4]. Therapeutic strategies that reverse these metabolic alterations while simultaneously targeting cancer-specific vulnerabilities remain an unmet clinical need.

Here we used time-series transcriptomics to characterize how crocin affects HCC cells over time. We examined whether crocin affects spliceosomal machinery and metabolic pathways implicated in HCC progression, and whether the two doses differ in pathway prioritization.

We employed a high-dose perturbation strategy (1–2 mM) to maximize transcriptional signal for mechanistic pathway discovery. This perturbation biology approach uses elevated concentrations to generate robust gene expression signatures that reveal drug-sensitive cellular machinery, a strategy employed in chemical genomics studies [11, 12]. By comparing two doses across multiple timepoints, we aimed to distinguish specific pathway effects from generalized stress responses and to identify candidate pathways that warrant investigation at physiologically relevant concentrations.

## 2 Results

### 2.1 Time-Series Transcriptomics Reveals Dynamic Cellular Response to Crocin

We performed comprehensive transcriptomic analysis of HepG2 cells treated with crocin at multiple timepoints (2, 6, 12, and 24 hours post-treatment (HPT)) using two concentrations (1 and 2 mM, corresponding to CR1 and CR2 treatments). The analysis employed transcription factor motif enrichment, gene ontology (GO) enrichment, and pathway analysis to characterize the cellular response.

Differential expression analysis identified substantial transcriptomic changes across all conditions (Table 1; full results in Table S2). Genes with | log_2_ FC| ≥ 1 and FDR-adjusted *p <* 0.05 were considered differentially expressed. For CR1, we observed 10,305 DEGs at 2 HPT (5960 upregulated, 4345 downregulated), decreasing to 7682 at 6 HPT and 7423 at 12 HPT, then rising to 9209 at 24 HPT. CR2 treatment yielded consistently higher total DEG counts: 11,329 at 2 HPT, 11,655 at 6 HPT, 12,076 at 12 HPT, and 10,186 at 24 HPT. CR2 showed a pronounced bias toward downregulation (7711 genes downregulated vs. 3618 upregulated at 2 HPT), while CR1 showed more balanced directionality.

**Table 1:** Differentially expressed gene counts by treatment condition and timepoint. DEGs were defined as genes with | log_2_ FC| ≥ 1 and FDR *<* 0.05 compared to T0 controls.

| Timepoint | CR1 Up | CR1 Down | CR1 Total | CR2 Up | CR2 Down | CR2 Total |
| --- | --- | --- | --- | --- | --- | --- |
| 2 HPT | 5960 | 4345 | 10,305 | 3618 | 7711 | 11,329 |
| 6 HPT | 2806 | 4876 | 7682 | 3631 | 8024 | 11,655 |
| 12 HPT | 2472 | 4951 | 7423 | 3729 | 8347 | 12,076 |
| 24 HPT | 5059 | 4150 | 9209 | 2950 | 7236 | 10,186 |

Despite CR2 producing 31% more total DEGs across all timepoints (45,246 vs. 34,619), pathway analysis revealed that CR1 achieved differential pathway prioritization on key cancer- relevant pathways (see Section 2.2). Gene overlap analysis showed moderate concordance between treatments, with Jaccard similarity indices ranging from 0.24 to 0.34 for cross-treatment comparisons at matched timepoints. Notably, CR2 DEG sets showed high internal consistency across timepoints (Jaccard 0.80–0.89), while CR1 showed greater temporal variation (Jaccard 0.24–0.58), indicating that higher-dose treatment produces a more uniform transcriptional response.

The response was complex and time-dependent: rapid, sustained downregulation of spliceosomal machinery coupled with activation of senescence and autophagy programs and suppression of proliferative pathways (Figure 1). HCC cells typically maintain persistent proliferative phenotypes with senescence programs suppressed [13], yet crocin disrupted this stability and activated senescence pathways.

**Figure 1.**
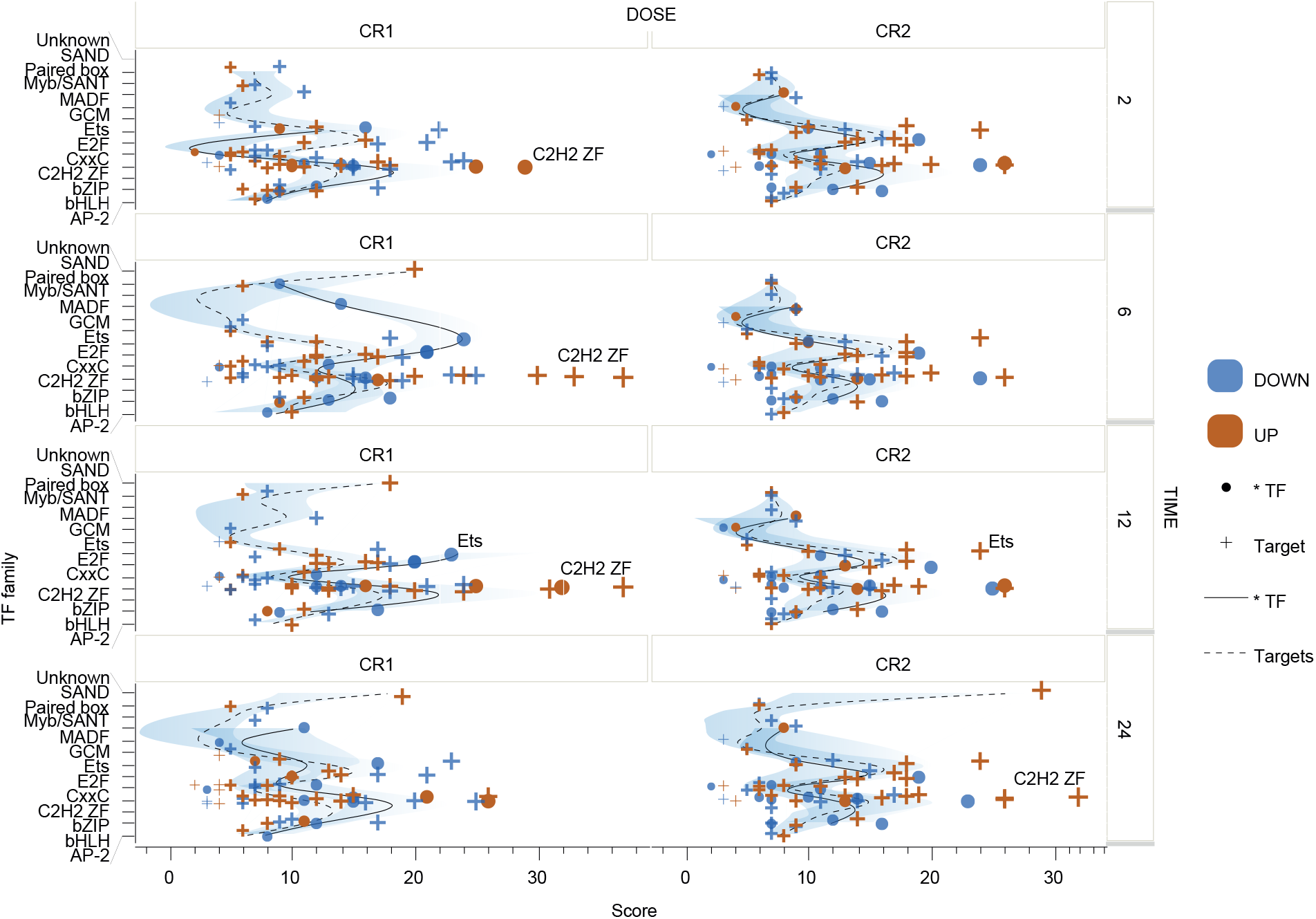
Transcription factor (TF) motif enrichment in differentially expressed genes following crocin treatment. Data: gene-level expression from RNA-seq; x-axis shows motif enrichment scores. Each row represents a timepoint (2, 6, 12, 24 HPT); columns show CR1 and CR2 treatments. TF families (y-axis) are plotted against enrichment score (x-axis). Blue symbols indicate enrichment among downregulated genes; orange symbols indicate enrichment among upregulated genes. Circles mark the TF itself; crosses mark TF target genes. Solid lines trace TF enrichment trends; dashed lines trace target gene enrichment trends across TF families. C2H2 zinc finger (C2H2 ZF) and Ets family TFs show consistently high enrichment scores. Genes containing promoter motifs for SP1/2, RREB1, PLAG1, PAX5, OSR1, EGR1, and ELF1 were rapidly upregulated, while ELK1 target genes showed decreased expression at 2 and 6 HPT.

### 2.2 Crocin Induces Progressive Downregulation of Spliceosomal Machinery

Spliceosomal components were consistently downregulated across all timepoints. Pathway enrichment analysis (hypergeometric test with Benjamini–Hochberg FDR correction; pathways with FDR *<* 0.05 considered significant; Table S5) revealed that the spliceosome was the top-ranked downregulated pathway for CR1 treatment at 2 HPT (FDR = 5.78 × 10^−21^, 72 genes), 6 HPT (FDR = 1.46 × 10^−36^, 93 genes), and 24 HPT (FDR = 6.06 × 10^−26^, 88 genes). In contrast, spliceosome ranked only fourth for CR2 at all timepoints (FDR = 10^−13^ to 10^−15^), despite CR2 affecting more total genes. This differential ranking indicates that lower-dose CR1 treatment achieves stronger enrichment for spliceosome pathways. This difference was reflected in pathway rankings: spliceosome ranked first among downregulated pathways for CR1 at three of four timepoints (mean rank 1.25) compared to fourth for CR2 at all timepoints (mean rank 4.0).

At 2 h, key affected genes included SRSF2 [14], RBM39, PRPF6, and SNRPC, affecting both core components and regulatory factors. By 6 HPT, this expanded to include additional SR proteins (SRSF7, SRSF8), their kinase SRPK1, and mRNA export factors (ALYREF, DDX39A). Expression analysis confirmed progressive downregulation, with DDX39B showing 37% reduction at 2 HPT and 62% reduction by 24 HPT in CR2-treated cells compared to controls.

This temporal progression from focused to broad spliceosome disruption suggests initial effects on key regulatory nodes followed by comprehensive machinery dysfunction. At 24 HPT, while some early changes persisted, new genes emerged (DDX39B, TXNL4A) and RNA degradation factors (EXOSC7) were affected, indicating establishment of a new RNA processing steady state (Figure 2).

**Figure 2.**
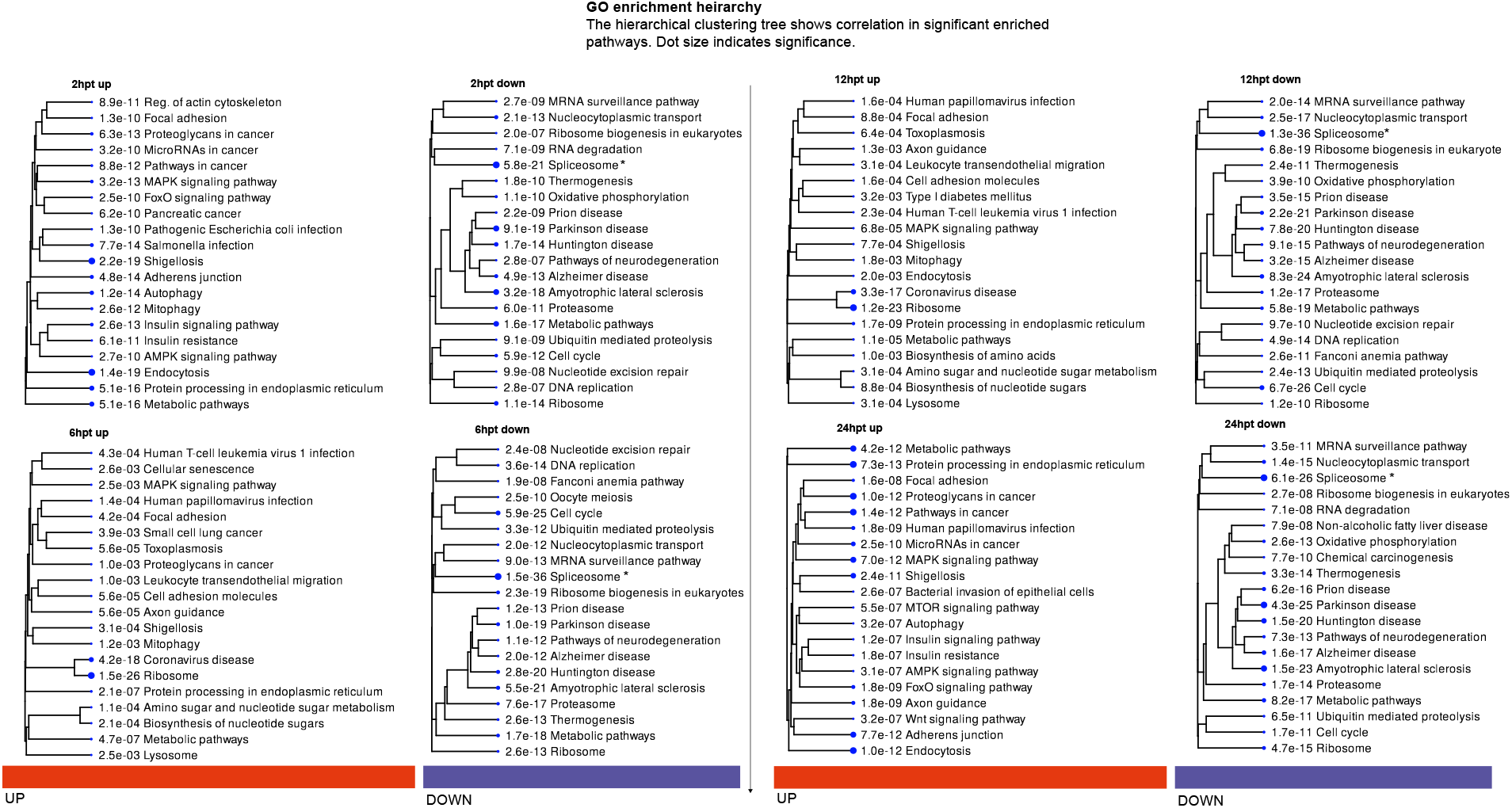
Gene ontology (GO) and Kyoto Encyclopedia of Genes and Genomes (KEGG) pathway enrichment hierarchies for CR1 treatment. Data: gene-level expression from RNA-seq; dot size indicates significance (FDR *p*-values shown); *n* = 7400–10,300 DEGs per timepoint. Eight panels show enriched pathways among upregulated (orange bar, left columns) and downregulated (blue bar, right columns) genes at each timepoint (2, 6, 12, 24 HPT). Hierarchical clustering trees group pathways by functional similarity. Spliceosome (marked with asterisks) consistently ranks among the top downregulated pathways across all timepoints (FDR 5.8 × 10^−21^ at 2 HPT to 1.3 × 10^−36^ at 6 HPT). Upregulated pathways include autophagy, cellular senescence, and MAPK signaling. Non-alcoholic fatty liver disease appears among downregulated pathways at 24 HPT.

### 2.3 Differential Splicing Analysis Validates Spliceosome Disruption

To determine whether the observed spliceosomal gene downregulation translates into functional splicing defects, we performed differential splicing analysis using SUPPA2 [15]. We analyzed 43,038 skipping exon (SE) events across all conditions (Tables S3, S4), comparing each treatment timepoint to T0 controls using the empirical method with 1000 bootstrap iterations to compute *p*-values. Events with |ΔPSI| ≥ 0.1 (representing ≥10 percentage point change in exon inclusion) and *p <* 0.05 were considered significant. This analysis revealed widespread splicing alterations, with 2620 significant events at CR1 2 HPT, decreasing to 584–692 at intermediate timepoints (6–12 HPT), then rising to 2072 at 24 HPT (Figure 3A). CR2 showed a similar biphasic pattern with 1748 events at 2 HPT and 2173 at 24 HPT.

**Figure 3.**
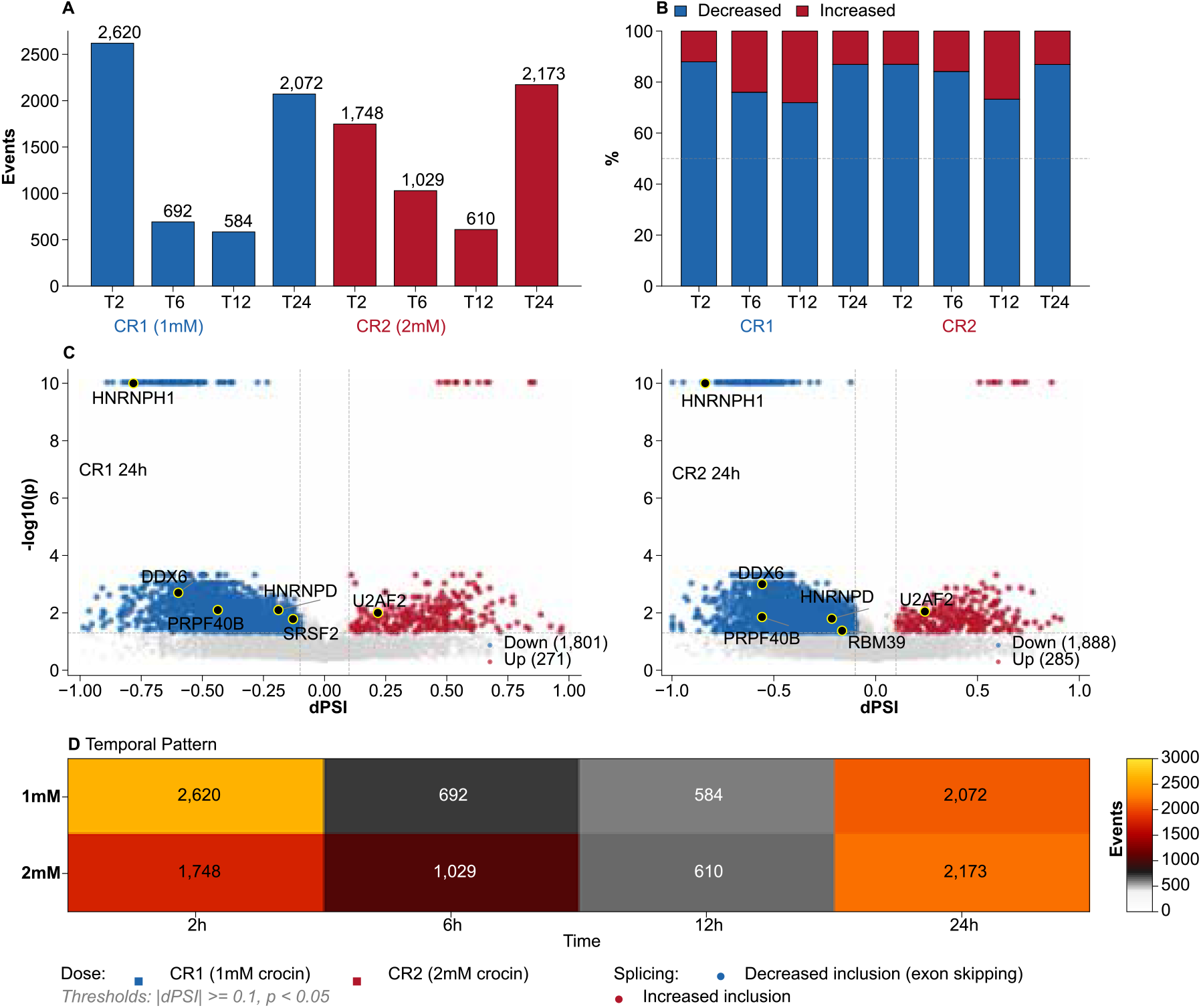
Differential splicing analysis of crocin-treated HCC cells. Data: transcript-level expression from RNA-seq; *n* = 43,038 SE events analyzed per condition. (**A**) Number of significant skipping exon (SE) events (|dPSI| ≥ 0.1, *p <* 0.05) at each timepoint for CR1 (1 mM) and CR2 (2 mM) treatment; *n* = 584−2620 significant events per condition. A biphasic pattern is evident with peaks at 2 HPT and 24 HPT. (**B**) Direction of splicing changes showing predominant exon skipping (decreased inclusion) across all conditions, with 72−88% of significant events showing negative dPSI values. (**C**) Volcano plots of differential splicing at 24 HPT for CR1 (left) and CR2 (right), with spliceosome component genes labeled. Blue points indicate decreased exon inclusion (exon skipping); red points indicate increased inclusion. Vertical dashed lines mark the |dPSI| = 0.1 significance threshold; y-axis shows − log_10_(*p*). (**D**) Temporal heatmap showing the number of significant splicing events across timepoints for both doses, illustrating the biphasic response pattern.

Across all conditions, 72–88% of significant splicing changes showed decreased exon inclusion (Figure 3B), a directional bias consistent with impaired splicing machinery. Among significant events, the median |ΔPSI| ranged from 0.21 to 0.42 across conditions (mean 0.31), with interquartile ranges of 0.15–0.29, indicating substantial effect magnitudes well above the 0.1 significance threshold.

Spliceosome component genes themselves showed altered splicing patterns (Figure 4). HN- RNPH1 [16], encoding heterogeneous nuclear ribonucleoprotein H1, exhibited dramatic exon skipping at a specific internal exon (chr5:179618145–179618323, 179 bp) with dPSI values of −0.78 to −0.89 across conditions (*p <* 0.001). This exon shows high baseline inclusion in control samples (mean percent spliced in, PSI = 0.95), indicating that crocin induces near-complete skipping of a normally constitutively included exon. This 179-bp cassette exon (exon 6 of the canonical ENST00000393432 transcript) resides entirely within the coding sequence, and its length (not divisible by 3) introduces a +2 frameshift upon skipping. The resulting premature termination codon is predicted to trigger nonsense-mediated decay (NMD), as seven exon-exon junctions remain downstream, placing the PTC well beyond the 50–55 nucleotide threshold from the last exon junction complex. Consequently, crocin treatment likely causes functional knockdown of HNRNPH1 through NMD-mediated transcript degradation. Given HNRNPH1’s central role in regulating alternative splicing of G-rich sequences, its loss may contribute to the broader splicing dysregulation observed in crocin-treated cells. Other affected spliceosome genes included PRPF40B [17] (dPSI = −0.44 to −0.56), DDX6 [18] (dPSI = −0.56 to −0.70), U2AF2 [19] (dPSI = +0.22 to +0.24, increased inclusion), and RBM39 [20] (dPSI = −0.16 to −0.24). The concordance between expression downregulation and aberrant splicing of spliceosome genes (Table 2) is consistent with a potential feedback model: crocin-induced expression changes impair splicing machinery, which in turn generates aberrantly spliced transcripts of spliceosome components, amplifying the dysfunction.

**Table 2:**
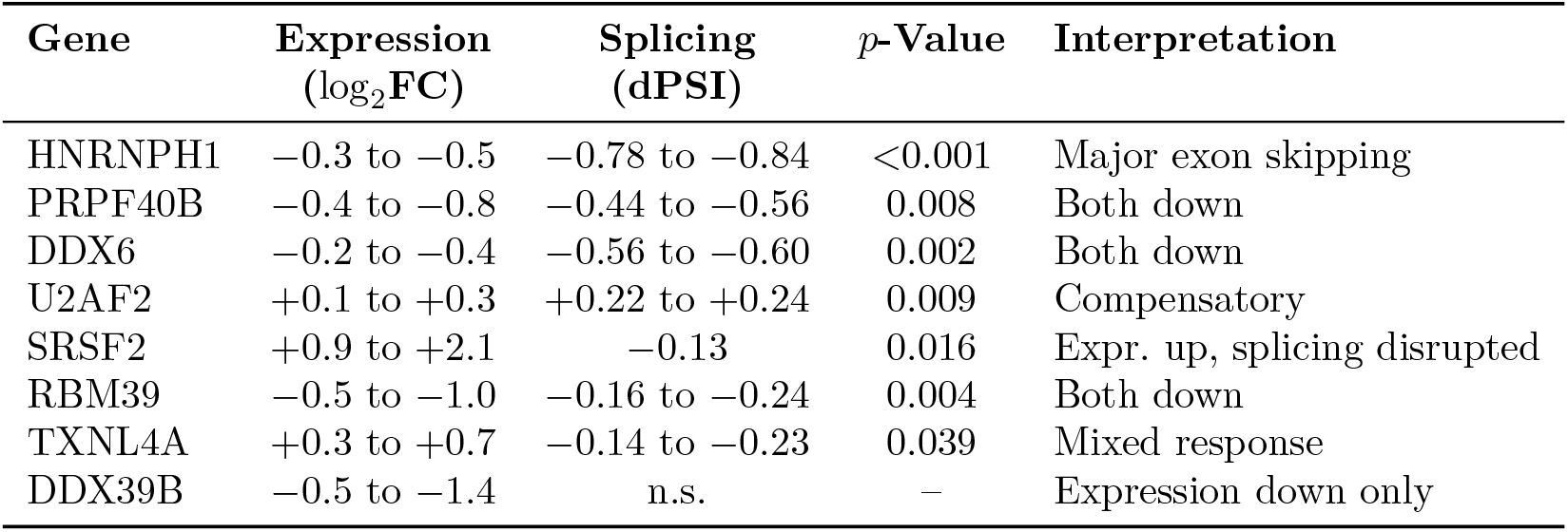
Spliceosome gene expression and splicing changes at 24 HPT. dPSI = change in percent spliced in; *p*-values from SUPPA2 empirical test. Single dPSI values indicate significance in one treatment condition only.

| Gene | Expression<br>(log<sub>2</sub>FC) | Splicing<br>(dPSI) | *p*-Value | Interpretation |
| --- | --- | --- | --- | --- |
| HNRNPH1 | −0.3 to −0.5 | −0.78 to −0.84 | <0.001 | Major exon skipping |
| PRPF40B | −0.4 to −0.8 | −0.44 to −0.56 | 0.008 | Both down |
| DDX6 | −0.2 to −0.4 | −0.56 to −0.60 | 0.002 | Both down |
| U2AF2 | +0.1 to +0.3 | +0.22 to +0.24 | 0.009 | Compensatory |
| SRSF2 | +0.9 to +2.1 | −0.13 | 0.016 | Expr. up, splicing disrupted |
| RBM39 | −0.5 to −1.0 | −0.16 to −0.24 | 0.004 | Both down |
| TXNL4A | +0.3 to +0.7 | −0.14 to −0.23 | 0.039 | Mixed response |
| DDX39B | −0.5 to −1.4 | n.s. | – | Expression down only |

**Figure 4.**
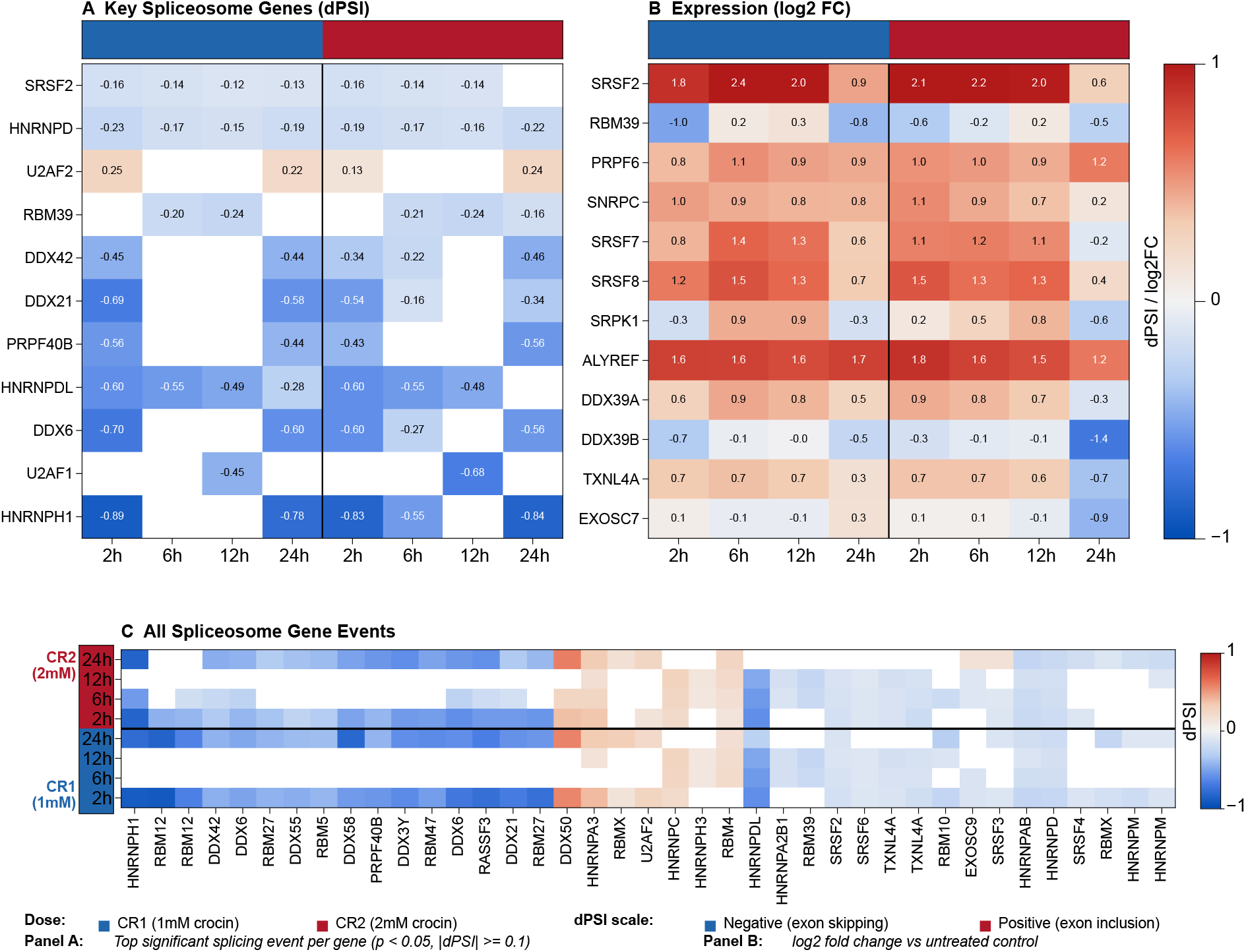
Differential splicing and expression of spliceosome component genes. Data: transcript- level (panels A, C) and gene-level (panel B) expression from RNA-seq. (**A**) dPSI values (x- axis) for key spliceosome genes (*n* = 8 genes shown) showing the most significant splicing event per gene (*p <* 0.05, |dPSI| ≥ 0.1). HNRNPH1 shows the most dramatic exon skipping (dPSI = −0.78 to −0.89), while U2AF2 uniquely shows increased exon inclusion (dPSI = +0.22 to +0.24). (**B**) Corresponding gene expression changes (log_2_ fold-change vs. untreated control) for key spliceosome genes affected at the splicing level or discussed in the text. (**C**) Comprehensive heatmap of all significant splicing events in spliceosome-associated genes across timepoints (*n* = 19 genes with defined SE events), showing the breadth of splicing disruption. Blue indicates decreased exon inclusion (exon skipping); red indicates increased inclusion. Gray cells indicate non-significant events at that timepoint.

### 2.4 Transcription Factor Analysis Reveals Chromatin Remodeling and Cell Fate Changes

Transcription factor motif enrichment analysis revealed distinct regulatory patterns governing crocin’s transcriptional response (Figure 1). Among C2H2 zinc finger transcription factors, SP1 [21] binding motifs showed the highest enrichment scores in upregulated genes across all timepoints (scores 25–31 for CR1), with SP2 motifs particularly enriched at 6 and 12 HPT (scores 37). EGR1 [22] (Early Growth Response 1) motifs were prominently enriched in upregulated genes (scores 26–33), peaking at 6 HPT, consistent with EGR1’s role as an immediate-early gene activated by growth factors and stress signals. PLAG1 and RREB1 motifs showed sustained enrichment in upregulated genes through 12 HPT (scores 16–24 and 10–17, respectively), suggesting coordinated activation of zinc finger-mediated transcriptional programs.

Notably, ELK1 [23] target genes showed an asymmetric pattern with preferential enrichment in downregulated genes (scores 16–18) compared to upregulated genes (scores 7–9) at 2 and 6 HPT for CR1 treatment. ELK1 regulates genes involved in redox homeostasis through glutathione metabolism pathways, and its target gene suppression suggests that crocin’s antioxidant properties may disrupt ELK1-dependent oncogenic signaling. The ETS family member ELF1 showed enrichment specifically in CR2-upregulated genes at 2, 6, and 24 HPT (scores 9–10), indicating dose-specific regulatory mechanisms.

PAX5 motifs presented a complex temporal pattern: while enriched in upregulated genes at early timepoints, PAX5 targets showed increased enrichment in downregulated gene sets at 24 HPT and across CR2 conditions. Given PAX5’s role in chromatin remodeling and lineage commitment, this shift may reflect epigenetic reprogramming as cells transition toward senescence-associated gene expression patterns observed at later timepoints.

### 2.5 Activation of Autophagy and Senescence Transcriptional Programs

KEGG pathway analysis revealed coordinated activation of the transcriptional signature of autophagy beginning at early timepoints. The autophagy pathway showed significant enrichment among upregulated genes at 12 HPT (21 genes, FDR = 0.051), including core autophagosome formation components (ATG5 [24], ATG12 [25], MAP1LC3B [26], MAP1LC3B2, GABARAPL1), WIPI proteins involved in autophagosome nucleation (WIPI1, WIPI2), and regulatory factors (BNIP3 [27], DDIT4 [28], DEPTOR). The RIG-I-like receptor signaling pathway, which interfaces with autophagy through ATG5 and ATG12, was also significantly enriched at 2, 12, and 24 HPT among upregulated genes.

Cellular senescence pathway genes showed dynamic regulation across timepoints. At 12 HPT, 26 senescence-associated genes were significantly upregulated (FDR = 0.011), including the key tumor suppressors CDKN2A [29] (p16^INK4a^)and CDKN1A [30] (p21), DNA damage response genes GADD45A and GADD45B [31], and senescence-associated secretory phenotype (SASP) components such as SERPINE1 [32], IL1A, and CXCL8. The p53 signaling pathway was concordantly activated with 14 upregulated genes at 12 HPT (FDR = 0.029), including BBC3/PUMA [33], TNFRSF10B, PMAIP1/NOXA [34], and SESN2 [35].

Simultaneously, cell cycle-promoting genes within the senescence pathway showed suppression: 46 genes were downregulated at 6 HPT (FDR = 0.004) including cyclins (CCND1, CCNE1, CCNB1/B2), CDKs (CDK2, CDK4, CDK6), and E2F transcription factors (E2F1, E2F2, E2F3, E2F4). This biphasic pattern—senescence activators up, proliferation genes down—was maintained through 24 HPT.

The temporal pattern of activation of senescence-associated transcriptional programs paralleled spliceosome disruption, raising the possibility of mechanistic connection. Spliceosome dysfunction acts as a gatekeeper for senescence entry, with pharmacologic or genetic perturbation of splicing factors sufficient to trigger transcriptional signatures of premature senescence [36]. At the molecular level, splicing defects promote R-loop accumulation and replication stress, activating ATR-mediated DNA damage responses that can lead to senescence-associated gene expression [37]. The mitophagy pathway (17 genes enriched at 12 HPT, FDR = 0.002), including PINK1 [38], BNIP3, BNIP3L, and BCL2L13, suggests coordinated organelle quality control accompanying the senescence transition. By 24 HPT, the senescence program appeared fully established, with sustained CDKN2A expression, cell cycle suppression, and SASP activation associated with transcriptional signatures of growth arrest without classical apoptosis.

### 2.6 Crocin Suppresses the Transcriptional Signature of NAFLD-Associated Metabolic Pathways

At 24 HPT, 66 genes associated with the NAFLD transcriptional signature were significantly downregulated (FDR *p* = 8 × 10^−8^, 1.96-fold enrichment). These included mitochondrial respiratory chain components—28 complex I subunits (NDUFA/B/C/S/V families) and ubiquinol-cytochrome c reductase subunits—along with cytochrome c oxidase subunits (COX6A1, COX6B1, COX6C, COX7A2, COX7C, COX8A) and metabolic signaling components (AKT1/2, MAPK9/11, PIK3CB, PPARG [39], SREBF1 [40], ADIPOR1/2).

Given that NAFLD progression to NASH represents a major HCC risk pathway, the coordinated suppression of these genes identifies metabolic pathways as crocin-sensitive under these experimental conditions. The downregulation of SREBF1 and PPARG, key regulators of lipogenesis, alongside adiponectin receptors indicates broad metabolic changes accompanying the anti-proliferative response. These metabolic gene changes could represent generalized metabolic suppression or cellular stress responses rather than NAFLD-specific pathway effects. The down- regulation of mitochondrial respiratory chain components and metabolic regulators is observed broadly in cellular stress and toxicity models, and therefore their potential therapeutic relevance for NAFLD-associated HCC requires validation at physiologically achievable concentrations in appropriate disease models.

### 2.7 Pathway Analysis Reveals Senescence-Associated Transcriptional Signatures Without Apoptosis

Gene Ontology analysis revealed time-dependent shifts: early responses (2–6 HPT) showed metabolic changes, while later timepoints featured sustained autophagy upregulation and continued spliceosome suppression. Ingenuity Pathway Analysis (IPA) at 24 HPT identified perturbations in molecular mechanisms of cancer, protein ubiquitination, and unfolded protein response pathways.

The unfolded protein response (UPR) pathway showed coordinated changes indicating ER stress adaptation rather than apoptotic commitment (Figure 5). PERK-mediated translational attenuation was active (EIF2*α* phosphorylation pathway engaged), while the IRE1-XBP1 [41] and ATF6 [42] branches showed mixed regulation. Downstream, CHOP-mediated BCL2 [43] activation promoted cell survival over apoptosis. The SREBP/PPARG axis showed downregulation, consistent with reduced lipogenesis observed in the NAFLD gene set. Heat shock proteins (HSPH1, HSP70, HSP40) were upregulated, indicating activation of protein quality control mechanisms.

**Figure 5.**
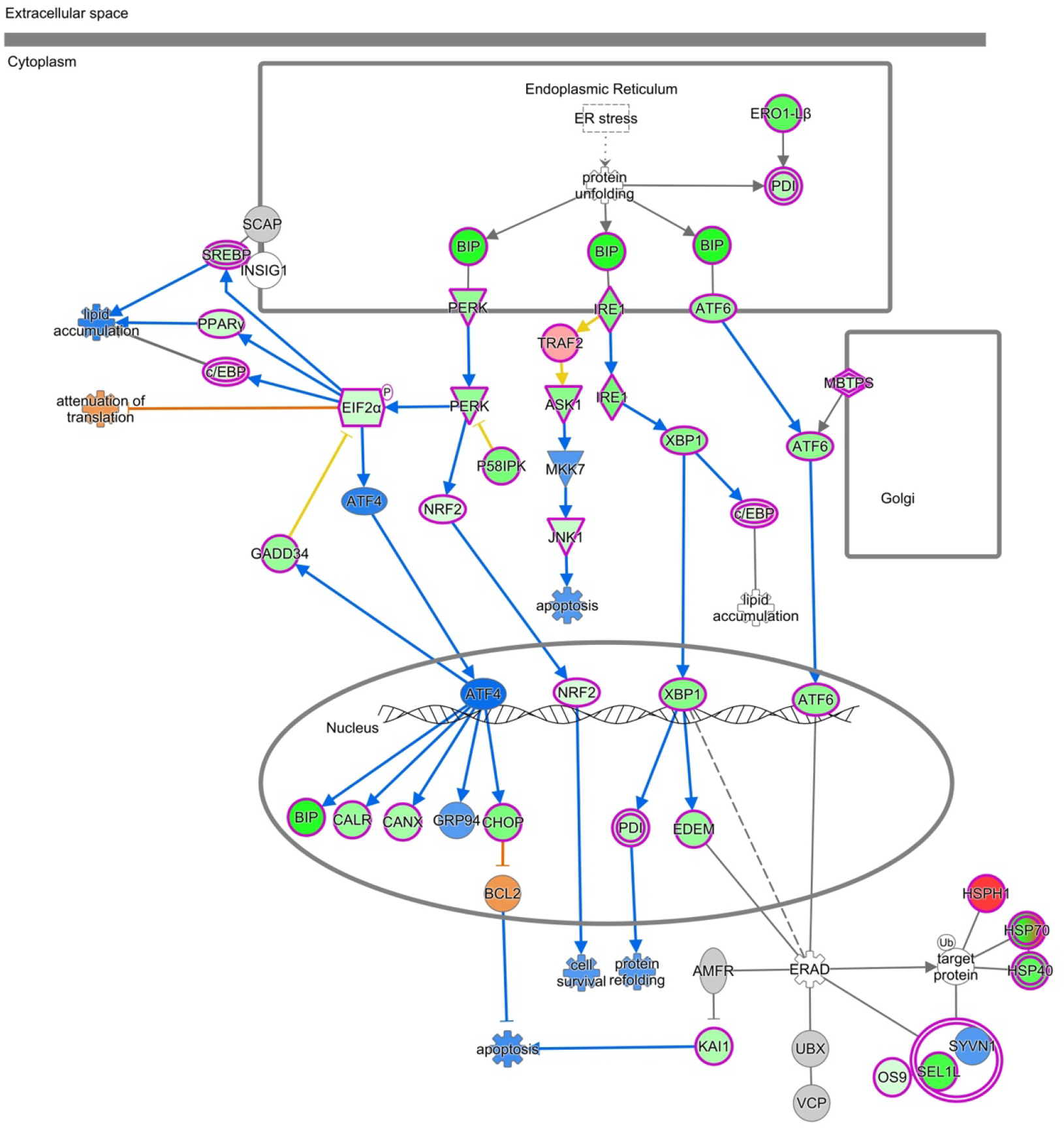
Ingenuity Pathway Analysis of the unfolded protein response (UPR) at 24 HPT. Data: gene-level expression from RNA-seq; node color intensity reflects log_2_ fold-change magnitude. Pathway map shows cellular compartments (ER, cytoplasm, nucleus, Golgi) with the three canonical UPR branches: PERK (left), IRE1 (center), and ATF6 (right). Node colors indicate expression changes: green = downregulated; red/pink = upregulated; gray = no significant change. Node shapes follow IPA conventions: rectangles = enzymes/kinases; ovals = transcription factors; diamonds = enzymes; trapezoids = transporters. CHOP-mediated BCL2 activation (bottom left) promotes cell survival over apoptosis. The SREBP/PPARG/cEBP lipogenesis axis (upper left) shows suppression. Heat shock proteins (HSPH1, HSP70, HSP40; lower right) are upregulated, indicating active protein quality control.

NF-*κ*B signaling was broadly suppressed (Figure 6). Multiple upstream activators (IL- 1R/TLR, TNFR, growth factor receptors) converge on the IKK complex, which showed reduced activation. Nuclear NF-*κ*B (p65/RelA, p50/p52) translocation was attenuated, with downstream consequences including reduced inflammatory gene expression, suppressed cell survival signaling, and decreased proliferation. The ASK1-MKK7-JNK1 pro-apoptotic arm was also downregulated, reinforcing the senescence-over-apoptosis phenotype.

**Figure 6.**
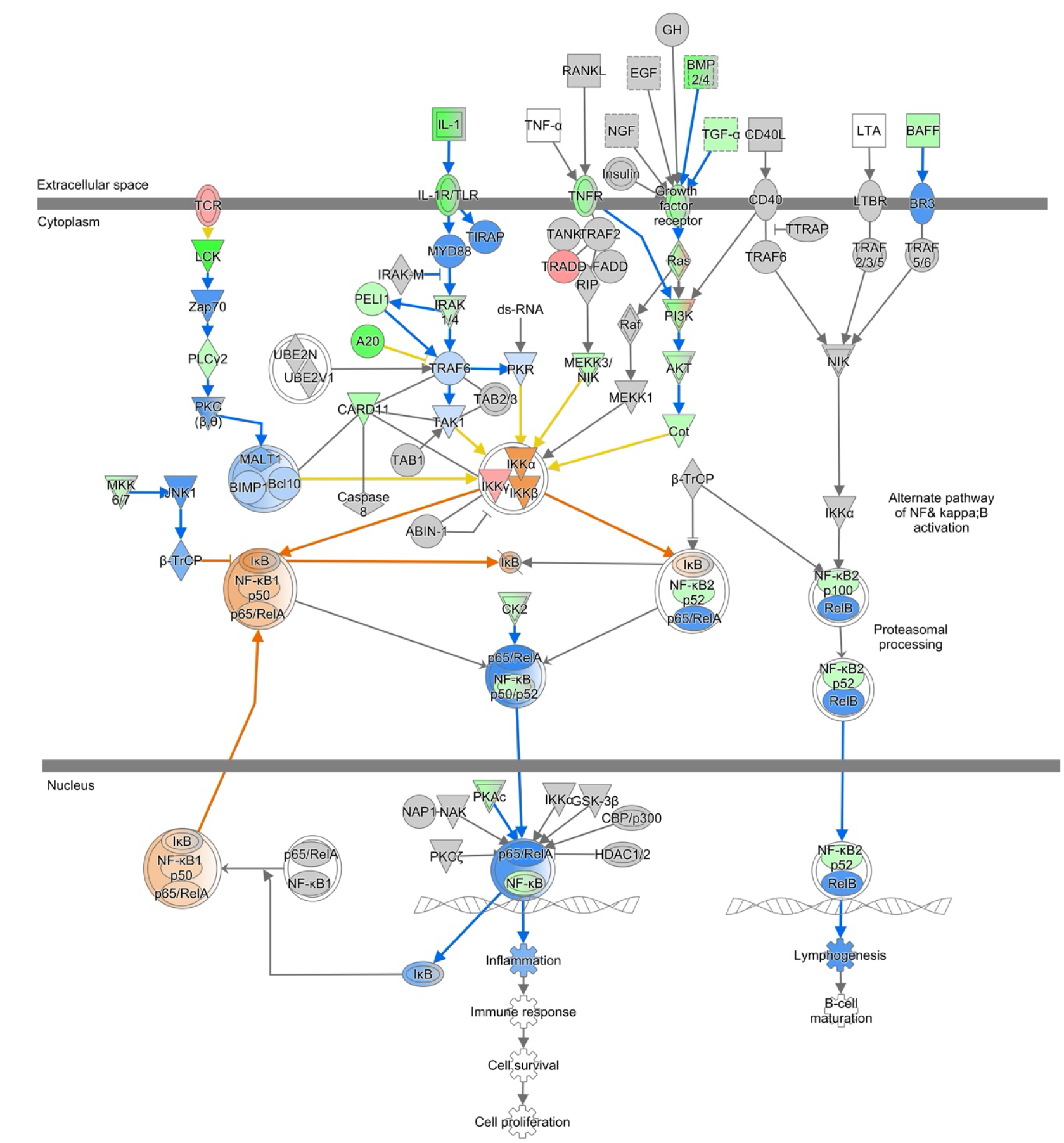
Ingenuity Pathway Analysis of NF-*κ*B signaling at 24 HPT. Data: gene-level expression from RNA-seq; node color intensity reflects log_2_ fold-change magnitude. Pathway map spans extracellular space (top), cytoplasm (middle), and nucleus (bottom). Multiple receptor inputs (IL-1, TNF-*α*, RANKL, EGF, growth factors) converge on the IKK complex (IKK*α*/IKK*β*) via adaptor proteins. Node colors indicate expression changes: green = downregulated; orange/red = upregulated; gray = no significant change. The classical NF-*κ*B pathway (left) shows I*κ*B degradation enabling p65/RelA and p50 nuclear translocation, while the alternative pathway (right) involves p100/RelB processing. Downstream transcriptional outputs (bottom) include inflammation, immune response, cell survival, and cell proliferation—all showing reduced activity. The MKK6/7-JNK1 pro-apoptotic arm (left side) is also suppressed, consistent with senescence rather than apoptosis.

## 3 Discussion

Using supraphysiological concentrations to probe maximal pathway engagement, our time-series analysis identifies the spliceosome as a primary pathway affected by crocin-induced cellular stress. Splicing dysregulation drives oncogenesis in *>*90% of cancers [6], and FDA-approved spliceosome modulators show efficacy in hematologic malignancies [8]. While spliceosome dysregulation in HCC has been documented [7], our findings reveal a distinct temporal pattern of disruption under crocin exposure. Alternative splicing events occur at *>*59-fold higher frequency than somatic mutations in HCC and generate substantially more immunogenic peptides [44]. Unlike the competing endogenous RNA networks that maintain splicing homeostasis in HCC [3, 4], crocin systematically perturbs this machinery.

Our SUPPA2 analysis provides direct functional validation, demonstrating that the predominant direction of splicing changes—decreased exon inclusion—is consistent with impaired splicing machinery. Spliceosome component genes themselves exhibited altered splicing, consistent with a potential feedback model: crocin-induced expression changes impair splicing machinery, which generates aberrantly spliced transcripts of spliceosome components, amplifying the dysfunction. HNRNPH1, which regulates splicing of numerous cancer-relevant transcripts [45, 46], showed near-complete exon skipping; Wen et al. demonstrated that HNRNPH1 competitively regulates PRMT5 splicing in HCC cells under radiation stress [45]. The differential response of individual components—some showing increased inclusion, others decreased—suggests genespecific rather than global splicing inhibition [47].

Notably, while the higher dose produced more total DEGs, the lower dose achieved differential pathway prioritization on cancer-relevant pathways, with spliceosome consistently ranking as the top downregulated pathway. This has implications for therapeutic optimization, suggesting that efficacy should be evaluated by pathway enrichment rather than total gene counts. The subsequent shift toward senescence—with CDKN2A upregulation and proliferation pathway suppression—represents a potent tumor-suppressive mechanism [13]. Dual activation of autophagy and senescence without apoptosis distinguishes crocin’s mechanism from conventional cytotoxic agents. Autophagy may initially serve as a survival response but can lead to cell death when hyperactivated [48, 49], and NF-*κ*B pathway downregulation further supports a pro-senescence phenotype. The coordinated downregulation of NAFLD-associated genes identifies metabolic pathways as crocin-sensitive; since NAFLD/NASH progression represents a major pathway to HCC development, suppression of lipogenesis regulators SREBF1 and PPARG warrants investigation at physiologically relevant concentrations. The concurrent downregulation of spliceosomal machinery and NAFLD-associated metabolic regulators suggests a potential regulatory axis. Recent studies indicate that PPAR*γ* and SREBF1 isoforms are heavily regulated by alternative splicing; thus, the crocin-induced spliceosome disruption may directly impair the processing of these metabolic transcription factors, leading to the observed mitochondrial and lipogenic suppression. This places the metabolic shift downstream of the RNA processing defects, rather than as a parallel, unrelated event.

### Scope and Limitations

Several limitations should be considered when interpreting these findings. The crocin concentrations used (1-2 mM) exceed reported plasma levels [50], and therefore the observed transcriptional responses likely reflect maximal pathway perturbation rather than direct modeling of therapeutic exposure; future studies at physiologically relevant concentrations (0.110 M) and extended treatment durations will clarify dose-dependent effects. Because time-matched untreated controls were not included, some expression changes may reflect temporal culture variation, although prior reports indicate relatively limited baseline fluctuation in HepG2 cells under stable conditions [5], and the magnitude of differential expression observed here supports a predominantly treatment-driven signal. HepG2 cells originate from a pediatric liver tumor and exhibit metabolic characteristics distinct from adult hepatocellular carcinoma models; therefore, validation in additional HCC cell lines and primary hepatocytes would strengthen generalizability. The association between spliceosomal disruption and senescence signatures cannot be resolved from transcriptomic data alone, as senescence was inferred from transcriptional markers rather than direct phenotypic assays. Crocin stability in culture medium was not directly assessed, and potential metabolites may contribute to the observed responses. Finally, the moderate overlap between treatment groups suggests partially distinct response programs that warrant further investigation.

## 4 Materials and Methods

### 4.1 HCC Cell Culture

The cell line present in this study was obtained from Sigma-Aldrich (Merck KGaA, Darmstadt, Germany). Cells were cultured in RPMI 1640 medium (HyClone, Logan, UT, USA) supplemented with 10% fetal bovine serum (Sigma-Aldrich, Burlington, MA, USA) containing 1% of antibiotic/antimycotic solution (Grand Island Biological Company (GIBCO); Thermo- Fisher Scientific, Waltham, MA, USA). Cells were incubated at 37*◦*C in a humidified 5% CO_2_ atmosphere and sub-cultured every 3–5 days using 0.25% trypsin (HyClone, Logan, UT, USA).

### 4.2 Crocin Treatment

HepG2 cells were incubated with 1 mM (CR1) or 2 mM (CR2) of crocin (Sigma-Aldrich, Burlington, MA, USA; ≥95% purity by HPLC) for 2, 6, 12, and 24 h time courses. These concentrations exceed achievable plasma levels following oral administration and were selected to induce maximal pathway perturbation for mechanistic discovery rather than to model therapeutic exposure. The osmotic contribution of crocin at 1−2 mM (*<*2 mOsm) is negligible relative to culture medium osmolarity (∼300 mOsm). Crocin solutions were prepared fresh immediately before treatment and protected from light during handling. Cells were trypsinized and pelleted. Cell pellets were resuspended in RNAlater stabilization solution (Ambion, from Invitrogen by Thermo Fischer Scientific, Carlsbad, CA, USA) and stored at −80^◦^C. All experiments were carried out in triplicates. Six untreated samples were taken at time point 0 and constitute the controls.

### 4.3 RNA-Seq Libraries Construction and Sequencing

Total RNA was isolated from three biological replicates of crocin-treated and untreated samples using an RNeasy Mini Kit (Qiagen, Hilden, Germany) following the manufacturer’s instructions. The RNA-seq libraries were prepared using TruSeq RNA sample prep kit (Illumina, Inc., San Diego, CA, USA) following the manufacturer’s instructions. Briefly, TruSeq RNA sample prep kit converts the poly-A containing mRNA in total RNA into a cDNA library using poly-T oligo-attached magnetic bead selection. Following mRNA purification, the RNA is chemically fragmented prior to reverse transcription and cDNA generation. The fragmentation step yields an RNA-seq library that includes inserts ranging in size from approximately 100–400 bp. The average insert size in an Illumina TruSeq RNA sequencing library is approximately 200 bp. The cDNA fragments then go through an end repair process, the addition of a single ‘A’ base to the 3*′* end followed by ligation of the adapters. Then, the products are purified and enriched with PCR to create the final double-stranded cDNA libraries. Finally, library quality control and quantification were performed using a Bioanalyzer Chip DNA 1000 series II (Agilent, Santa Clara, CA, USA) and sequenced directly using the high-throughput Illumina HiSeq sequencing system (Illumina, Inc., San Diego, CA, USA).

### 4.4 Alignment and Analysis of Illumina Reads Against the Reference Genome

Paired End (PE) reads from each library (crocin treatments with selected time points and in addition to the control sample) were generated using CASAVA version 1.8.2 package in Fastq format. The Trimmomatic [51, 52] package was used to remove adapters and filter out low quality bases (Q *<* 20). Filtered sequences reads were mapped to the ENSEMBL [53] human genome sequence (ENSEMBL-release-81-GRCh38), which was used as a reference to align the transcriptome reads. To characterize gene expression in cells, TPM values (Transcripts Per Million) were calculated using Kallisto [54] and used as a normalized measure of transcript abundance that allows comparison both within and between samples. Kallisto pseudoalignment was performed against ENSEMBL transcriptome sequences (release 81, GRCh38) with default parameters (Table S1).

Differentially expressed genes (DEGs) were identified by comparing each treatment condition to T0 controls using Sleuth (v0.30.0) [55] with Kallisto abundance estimates. Genes were considered differentially expressed if they met two criteria: absolute log_2_ fold-change ≥ 1 and false discovery rate (FDR) adjusted *p*-value *<* 0.05. This threshold identified 7400−12,100 DEGs per condition. A separate analysis using DESeq2 [56] with count data from RSEM [57] was performed specifically for Ingenuity Pathway Analysis (IPA), using the same significance thresholds.

### 4.5 Ingenuity Pathway Analysis

We used the Ingenuity Pathway Analysis (IPA; Ingenuity Systems, Inc., Redwood City, CA, USA) to examine the biological network associated with changes in RNA expression levels from the crocin treatment at 12 and 24 HPT. The IPA software (http://www.ingenuity.com) used a manually curated database which contains information from several sources, including published journal papers and gene annotation databases. Fisher’s exact test was used to calculate the probabilities between the input gene set and the canonical pathways. We also used IPA to predict the upstream and downstream effects of activation or inhibition on other molecules based on the input gene set’s expression data.

### 4.6 GO Analyses

All query genes are first converted to ENSEMBL [53] gene IDs or STRING-db [58] protein IDs. Our gene ID mapping and pathway data were derived from these two sources and processed using ShinyGO (https://bioinformatics.sdstate.edu/go/) [59]. False discovery rate (FDR) was calculated based on the nominal *p*-value from the hypergeometric test. Fold-enrichment was defined as the percentage of genes in the list belonging to a pathway, divided by the corresponding percentage in the background. Due to increased statistical power, large pathways tend to have smaller FDRs. As a measure of effect size, fold-enrichment indicated the degree of overrepresentation of genes in a pathway. Only pathways that are within the specified size limits were used for enrichment analysis. Pathways were then filtered based on a user-specified FDR cutoff. Then the significant pathways were sorted by FDR, fold-enrichment, or other metrics. We first selected the top pathways by FDR, then sorted these by fold enrichment. Similar pathways sharing 95% of their genes were represented instead by the single most significant pathway. Redundant pathways were required to share 50% of the words in their names.

### 4.7 Statistical Analysis

Statistical analyses were performed using JMP software (version 16.0 PRO, SAS Institute Inc., Cary, NC, USA). For continuous dependent variables and categorical independent variables, we conducted pairwise comparisons across all levels of the categorical variables. Each comparison included a standard *t*-test, a test of practical significance, and an equivalence test. We implemented an outlier detection method for each continuous dependent variable and categorical independent variable. Outliers were defined as points whose distance to the predicted value exceeded three times the estimated sigma (i.e., points with standardized residuals exceeding three). These outlier indicators were saved as new columns in the original data table, grouped under the ‘outlier group’ category. Robust fits and robust estimates were used for outlier detection, with the same three-sigma criterion applied. For both residual analysis and outlier detection, we performed robustness checks by repeating the analyses with the robust option selected.

### 4.8 Differential Splicing Analysis

To quantify changes in alternative splicing, we performed differential splicing analysis using SUPPA2 [15]. Transcript-level TPM values from Kallisto quantification were used as input. Note that while Kallisto quantification used ENSEMBL transcriptome annotation (Section 4.4), splicing event definitions were generated from RefSeq annotation (GRCh38.p14) using the SUPPA2 generateEvents module with the --pool-genes option to aggregate transcripts by gene. Transcript IDs were mapped between annotation systems using gene symbols. Transcript mapping achieved 100% coverage (177,816 transcripts), with 19 of 24 key spliceosome genes analyzed in this study having defined SE events in the annotation. RefSeq annotation was selected for splicing analysis because its exon boundaries are more conservatively curated based on experimental evidence, which improves specificity for detecting biologically meaningful splicing changes [60]. The complete transcript mapping coverage (100%) and internal consistency of results across multiple timepoints support the validity of this approach.

We analyzed 43,038 skipping exon (SE) events, representing 31.6% of the 136,239 total alternative splicing events defined in our RefSeq annotation. SE events are the most prevalent and best-characterized class of alternative splicing [61] and are mechanistically interpretable, as they represent discrete inclusion/exclusion decisions for internal exons. Other event types— alternative first exons (36,698; 26.9%), alternative 5*′*/3*′* splice sites (34,856; 25.6%), and retained introns (6,659; 4.9%)—were not analyzed in this study but represent avenues for future investigation.

Percent spliced in (PSI) values were calculated for each condition using the psiPerEvent module. Differential splicing analysis was performed using the diffSplice module with the empirical method, comparing each treatment condition (CR1 and CR2 at 2, 6, 12, and 24 HPT) to T0 controls. Events were considered significantly differentially spliced if they met two criteria: absolute change in PSI (|dPSI|) ≥ 0.1 and empirical *p*-value *<* 0.05. The 0.1 dPSI threshold corresponds to a 10 percentage point change in exon inclusion and represents a biologically meaningful shift in splicing outcome.

To identify splicing changes in spliceosome component genes specifically, we extracted events mapping to genes annotated in the KEGG spliceosome pathway (hsa03040) and manually curated lists of splicing factors (SR proteins, hnRNPs, and core spliceosomal components).

## 5 Conclusions

Using supraphysiological crocin concentrations to probe maximal pathway engagement, our time-series transcriptomic analysis of HepG2 cells identifies spliceosome components as a primary crocin-sensitive pathway. Crocin exposure induced 7400–12,100 DEGs per condition, with the spliceosome emerging as the top downregulated pathway for CR1 treatment (FDR 10^−21^ to 10^−36^). The lower dose achieved differential pathway prioritization compared to the higher dose, consistent with dose-dependent transition from focused pathway enrichment to generalized stress responses. Additionally, 66 NAFLD-associated genes were downregulated at 24 HPT, and crocin induced transcriptional features consistent with senescence rather than apoptosis, indicating growth arrest as the primary cellular response.

These findings identify spliceosome components and RNA processing machinery as crocin- sensitive pathways. Our differential splicing analysis provides functional validation, demonstrating that crocin exposure not only downregulates spliceosome gene expression but also induces widespread aberrant splicing, including of spliceosome components themselves. However, the supraphysiological concentrations used here substantially exceed achievable tissue levels, and the observed effects should be interpreted as identifying candidate pathways rather than demonstrating therapeutic mechanisms. Future studies using physiologically relevant concentrations (0.1–10 *µ*M), extended exposure times, and multiple cell lines will be essential to determine whether the spliceosome sensitivity we identified translates to therapeutic contexts.

## Supplementary Materials

The following supporting information can be downloaded at: Zenodo doi:10.5281/zenodo.12632036 (https://zenodo.org/records/15804191): Table S1 (transcript-level TPM expression matrix), Table S2 (differential expression analysis results), Table S3 (PSI matrix for skipping exon events), Table S4 (differential splicing analysis results), Table S5 (KEGG pathway enrichment results), and raw sequencing data.

## Author Contributions

**D.R.N**.: software, validation, formal analysis, investigation, data curation, writing—review and editing, visualization. **A.C**.: software, validation, formal analysis, investigation, data curation, writing—original draft, writing—review and editing, visualization. **W.F**.: writing—review and editing. **A.S.A**.: formal analysis, investigation. **A.A.-H**.: formal analysis, investigation. **A.A**.: conceptualization, methodology, resources, writing—original draft, writing—review and editing, supervision, project administration, funding acquisition. **K.S.-A**.: conceptualization, methodology, resources, writing—original draft, writing—review and editing, supervision, project administration, funding acquisition. All authors have read and agreed to the published version of the manuscript.

The authors used Claude Opus 4.5 (Anthropic) to assist with manuscript editing. The authors reviewed and edited all output and take full responsibility for the content of this publication.

## Institutional Review Board Statement

Not applicable.

## Informed Consent Statement

Not applicable.

## Data Availability Statement

All data used for these analyses have been uploaded to online repositories with permanently accessible digital object identifiers (DOIs) to ensure reproducibility and facilitate further exploration of the results (Zenodo doi:10.5281/zenodo.12632036; https://zenodo.org/records/15804191). Sequencing reads are available at Zenodo doi:10.5281/zenodo.13822023 (controls; https://zenodo.org/records/13822024) and doi:10.5281/zenodo.13842438 (crocin treatment time points 2, 6, 12, and 24 HPT; https://zenodo.org/records/13842439).

## Abbreviations

The following abbreviations are used in this manuscript:

DEG: Differentially expressed gene
dPSI: Change in percent spliced in
FDR: False discovery rate
GO: Gene ontology
HCC: Hepatocellular carcinoma
HPT: Hours post treatment
IPA: Ingenuity Pathway Analysis
NAFLD: Non-alcoholic fatty liver disease
NASH: Non-alcoholic steatohepatitis
PSI: Percent spliced in
SE: Skipping exon
TF: Transcription factor
TPM: Transcripts per million

## References

[1] C. Z. Wang, Q. Ma, S. Kim, D. H. Wang, Y. Shoyama, and C. S. Yuan. Effects of saffron and its active constituent crocin on cancer management: A narrative review. Longhua Chin. Med., 5:35, 2022.

[2] B. Awad, A. A. Hamza, A. Al-Maktoum, S. Al-Salam, and A. Amin. Combining crocin and sorafenib improves their tumor-inhibiting effects in a rat model of diethylnitrosamine-induced cirrhotic-hepatocellular carcinoma. Cancers, 15:4063, 2023.

[3] Z. Tang, X. Li, Y. Zheng, J. Liu, C. Liu, and X. Li. The role of competing endogenous RNA network in the development of hepatocellular carcinoma: Potential therapeutic targets. Front. Cell Dev. Biol., 12:1341999, 2024.

[4] Z. S. Niu, W. H. Wang, X. N. Dong, and L. M. Tian. Role of long noncoding RNA-mediated competing endogenous RNA regulatory network in hepatocellular carcinoma. World J. Gastroenterol., 26:4240–4260, 2020.

[5] A. V. Tyakht, E. N. Ilina, D. G. Alexeev, D. S. Ischenko, A. Y. Gorbachev, T. A. Semashko, A. K. Larin, O. V. Selezneva, E. S. Kostryukova, P. A. Karalkin, et al. RNA-Seq gene expression profiling of HepG2 cells: The influence of experimental factors and comparison with liver tissue. BMC Genomics, 15:1108, 2014.

[6] L. M. Urbanski, N. Leclair, and O. Anczukow. Alternative-splicing defects in cancer: Splicing regulators and their downstream targets, guiding the way to novel cancer therapeutics. Wiley Interdiscip. Rev. RNA, 9:e1476, 2018.

[7] S. E. Lee, K. P. Alcedo, H. J. Kim, and N. T. Snider. Alternative splicing in hepatocellular carcinoma. Cell. Mol. Gastroenterol. Hepatol., 10:699–712, 2020.

[8] Q. Peng, Y. Zhou, L. Oyang, N. Wu, Y. Tang, M. Su, X. Luo, Y. Wang, X. Sheng, J. Ma, et al. Impacts and mechanisms of alternative mRNA splicing in cancer metabolism, immune response, and therapeutics. Mol. Ther., 30:1018–1035, 2022.

[9] X. Lv, X. Sun, Y. Gao, X. Song, X. Hu, L. Gong, L. Han, M. He, and M. Wei. Targeting RNA splicing modulation: New perspectives for anticancer strategy? J. Exp. Clin. Cancer Res., 44:21, 2025.

[10] S. Araki, M. Ohori, and M. Yugami. Targeting pre-mRNA splicing in cancers: Roles, inhibitors, and therapeutic opportunities. Front. Oncol., 13:1152087, 2023.

[11] J. Lamb, E. D. Crawford, D. Peck, J. W. Modell, I. C. Blat, M. J. Wrobel, J. Lerner, J. P. Brunet, A. Subramanian, K. N. Ross, M. Reich, H. Hieronymus, G. Wei, S. A. Armstrong, S. J. Haggarty, P. A. Clemons, R. Wei, S. A. Carr, E. S. Lander, and T. R. Golub. The connectivity map: Using gene-expression signatures to connect small molecules, genes, and disease. Science, 313:1929–1935, 2006.

[12] A. Subramanian, R. Narayan, S. M. Corsello, D. D. Peck, T. E. Natoli, X. Lu, J. Gould, J. F. Davis, A. A. Tubelli, J. K. Asiedu, D. L. Lahr, J. E. Hirschman, Z. Liu, M. Donahue, B. Julian, M. Khan, D. Wadden, I. C. Smith, D. Lam, A. Liberzon, et al. A next generation connectivity map: L1000 platform and the first 1,000,000 profiles. Cell, 171:1437–1452.e17, 2017.

[13] D. V. Faget, Q. Ren, and S. A. Stewart. Unmasking senescence: Context-dependent effects of SASP in cancer. Nat. Rev. Cancer, 19:439–453, 2019.

[14] Y. Komeno, Y. J. Huang, J. Qiu, L. Lin, Y. J. Xu, Y. Zhou, L. Chen, D. D. Monterroza, H. Li, R. C. DeKelver, et al. SRSF2 is essential for hematopoiesis, and its myelodysplastic syndrome-related mutations dysregulate alternative pre-mRNA splicing. Mol. Cell. Biol., 35:3071–3082, 2015.

[15] J. L. Trincado, J. C. Entizne, G. Hysenaj, B. Singh, M. Skaber, D. J. Elliott, and E. Eyras. SUPPA2: Fast, accurate, and uncertainty-aware differential splicing analysis across multiple conditions. Genome Biol., 19:40, 2018.

[16] L. Zhu, W. Yi, L. Zhang, C. Qiu, N. Sun, J. He, P. Feng, Q. Wu, G. Wang, and G. Wu. hnRNPH1: A multifaceted regulator in RNA processing and disease pathogenesis. Int. J. Mol. Sci., 26:5159, 2025.

[17] S. Becerra, M. Montes, C. Hernández-Munain, and C. Suñé. Prp40 pre-mRNA processing factor 40 homolog B (PRPF40B) associates with SF1 and U2AF65 and modulates alternative pre-mRNA splicing in vivo. RNA, 21:438–457, 2015.

[18] B. Di Stefano, E. C. Luo, C. Haggerty, S. Aigner, J. Charlton, J. Brumbaugh, F. Ji, I. Rabano Jiménez, K. J. Clowers, A. J. Huebner, et al. The RNA helicase DDX6 controls cellular plasticity by modulating P-body homeostasis. Cell Stem Cell, 25:622–638.e13, 2019.

[19] S. Wu, C. M. Romfo, T. W. Nilsen, and M. R. Green. Functional recognition of the 3’ splice site AG by the splicing factor U2AF35. Nature, 402:832–835, 1999.

[20] Y. Xu, A. Nijhuis, and H. C. Keun. RNA-binding motif protein 39 (RBM39): An emerging cancer target. Br. J. Pharmacol., 179:2795–2812, 2022.

[21] S. S. Solomon, G. Majumdar, A. Martinez-Hernandez, and R. Raghow. A critical role of Sp1 transcription factor in regulating gene expression in response to insulin and other hormones. Life Sci., 83:305–312, 2008.

[22] B. Wang, H. Guo, H. Yu, Y. Chen, H. Xu, and G. Zhao. The role of the transcription factor EGR1 in cancer. Front. Oncol., 11:642547, 2021.

[23] M. Patki, V. Chari, S. Sivakumaran, M. Gonit, R. Trumbly, and M. Ratnam. The ETS domain transcription factor ELK1 directs a critical component of growth signaling by the androgen receptor in prostate cancer cells. J. Biol. Chem., 288:11047–11065, 2013.

[24] X. Ye, X. J. Zhou, and H. Zhang. Exploring the role of autophagy-related gene 5 (ATG5) yields important insights into autophagy in autoimmune/autoinflammatory diseases. Front. Immunol., 9:2334, 2018.

[25] J. Geng and D. J. Klionsky. The Atg8 and Atg12 ubiquitin-like conjugation systems in macroautophagy. EMBO Rep., 9:859–864, 2008.

[26] I. Tanida, T. Ueno, and E. Kominami. LC3 and autophagy. Methods Mol. Biol., 445:77–88, 2008.

[27] J. Zhang and P. A. Ney. Role of BNIP3 and NIX in cell death, autophagy, and mitophagy. Cell Death Differ., 16:939–946, 2009.

[28] J. Brugarolas, K. Lei, R. L. Hurley, B. D. Manning, J. H. Reiling, E. Hafen, L. A. Witters, L. W. Ellisen, and W. G. Kaelin. Regulation of mTOR function in response to hypoxia by REDD1 and the TSC1/TSC2 tumor suppressor complex. Genes Dev., 18:2893–2904, 2004.

[29] J. P. Coppé, F. Rodier, C. K. Patil, A. Freund, P. Y. Desprez, and J. Campisi. Tumor suppressor and aging biomarker p16(INK4a) induces cellular senescence without the associated inflammatory secretory phenotype. J. Biol. Chem., 286:36396–36403, 2011.

[30] J. Yan, S. Chen, Z. Yi, R. Zhao, J. Zhu, S. Ding, and J. Wu. The role of p21 in cellular senescence and aging-related diseases. Mol. Cells, 47:100113, 2024.

[31] A. Humayun and A. J. Fornace. GADD45 in stress signaling, cell cycle control, and apoptosis. Adv. Exp. Med. Biol., 1360:1–22, 2022.

[32] D. E. Vaughan, R. Rai, S. S. Khan, M. Eren, and A. K. Ghosh. Plasminogen activator inhibitor-1 is a marker and a mediator of senescence. Arterioscler. Thromb. Vasc. Biol., 37:1446–1452, 2017.

[33] J. R. Jeffers, E. Parganas, Y. Lee, C. Yang, J. Wang, J. Brennan, K. H. MacLean, J. Han, T. Chittenden, J. N. Ihle, et al. Puma is an essential mediator of p53-dependent and -independent apoptotic pathways. Cancer Cell, 4:321–328, 2003.

[34] E. Oda, R. Ohki, H. Murasawa, J. Nemoto, T. Shibue, T. Yamashita, T. Tokino, T. Taniguchi, and N. Tanaka. Noxa, a BH3-only member of the Bcl-2 family and candidate mediator of p53-induced apoptosis. Science, 288:1053–1058, 2000.

[35] C. Lu, Y. Jiang, W. Xu, and X. Bao. Sestrin2: Multifaceted functions, molecular basis, and its implications in liver diseases. Cell Death Dis., 14:160, 2023.

[36] S. M. Kwon, S. Min, U. W. Jeoun, M. S. Sim, G. H. Jung, S. M. Hong, B. A. Jee, H. G. Woo, C. Lee, and G. Yoon. Global spliceosome activity regulates entry into cellular senescence. FASEB J., 35:e21204, 2021.

[37] H. D. Nguyen, W. Y. Leong, W. Li, P. N. G. Reddy, J. D. Sullivan, M. J. Walter, L. Zou, and T. A. Graubert. Spliceosome mutations induce R loop-associated sensitivity to ATR inhibition in myelodysplastic syndromes. Cancer Res., 78:5363–5374, 2018.

[38] A. M. Pickrell and R. J. Youle. The roles of PINK1, parkin, and mitochondrial fidelity in Parkinson’s disease. Neuron, 85:257–273, 2015.

[39] O. Gavrilova, M. Haluzik, K. Matsusue, J. J. Cutson, L. Johnson, K. R. Dietz, C. J. Nicol, C. Vinson, F. J. Gonzalez, and M. L. Reitman. Liver peroxisome proliferator-activated receptor gamma contributes to hepatic steatosis, triglyceride clearance, and regulation of body fat mass. J. Biol. Chem., 278:34268–34276, 2003.

[40] N. Li, X. Li, Y. Ding, X. Liu, K. Diggle, T. Kisseleva, and D. A. Brenner. SREBP regulation of lipid metabolism in liver disease, and therapeutic strategies. Biomedicines, 11:3280, 2023.

[41] H. Yoshida, T. Matsui, A. Yamamoto, T. Okada, and K. Mori. XBP1 mRNA is induced by ATF6 and spliced by IRE1 in response to ER stress to produce a highly active transcription factor. Cell, 107:881–891, 2001.

[42] K. Haze, H. Yoshida, H. Yanagi, T. Yura, and K. Mori. Mammalian transcription factor ATF6 is synthesized as a transmembrane protein and activated by proteolysis in response to endoplasmic reticulum stress. Mol. Biol. Cell, 10:3787–3799, 1999.

[43] S. Qian, Z. Wei, W. Yang, J. Huang, Y. Yang, and J. Wang. The role of BCL-2 family proteins in regulating apoptosis and cancer therapy. Front. Oncol., 12:985363, 2022.

[44] H. Zhao, Y. Cheng, T. Zhang, Q. Wang, Y. Xu, G. Wang, Y. Song, H. Chen, Y. Wu, M. Zhang, et al. Harnessing alternative splicing for off-the-shelf mRNA neoantigen vaccines in hepatocellular carcinoma. Cell Res., 35:970–986, 2025.

[45] C. Wen, Z. Tian, L. Li, T. Chen, H. Chen, J. Dai, Z. Liang, S. Ma, and X. Liu. SRSF3 and HNRNPH1 regulate radiation-induced alternative splicing of protein arginine methyl-transferase 5 in hepatocellular carcinoma. Int. J. Mol. Sci., 23:14832, 2022.

[46] T. Brownmiller and N. J. Caplen. The HNRNPF/H RNA binding proteins and disease. Wiley Interdiscip. Rev. RNA, 14:e1788, 2023.

[47] A. Bashari, Z. Siegfried, and R. Karni. Targeting splicing factors for cancer therapy. RNA, 29:506–515, 2023.

[48] A. M. Leidal, B. Levine, and J. Debnath. Autophagy and the cell biology of age-related disease. Nat. Cell Biol., 20:1338–1348, 2018.

[49] N. Mizushima and B. Levine. Autophagy in human diseases. N. Engl. J. Med., 383: 1564–1576, 2020.

[50] A. Asai, T. Nakano, M. Takeuchi, C. Murakami, and T. Miyazawa. Orally administered crocetin and crocins are absorbed into blood plasma as crocetin and its glucuronide conjugates in mice. J. Agric. Food Chem., 53:7302–7306, 2005.

[51] S. O. Sewe, G. Silva, P. Sicat, S. E. Seal, and P. Visendi. Trimming and validation of illumina short reads using trimmomatic, trinity assembly, and assessment of RNA-Seq data. Methods Mol. Biol., 2443:211–232, 2022.

[52] A. M. Bolger, M. Lohse, and B. Usadel. Trimmomatic: A flexible trimmer for illumina sequence data. Bioinformatics, 30:2114–2120, 2014.

[53] S. C. Dyer, O. Austine-Orimoloye, A. G. Azov, M. Barba, I. Barnes, V. P. Barrera-Enriquez, A. Becker, R. Bennett, M. Beracochea, A. Berry, et al. Ensembl 2025. Nucleic Acids Res., 53:D948–D957, 2025.

[54] N. L. Bray, H. Pimentel, P. Melsted, and L. Pachter. Near-optimal probabilistic RNA-seq quantification. Nat. Biotechnol., 34:525–527, 2016.

[55] H. Pimentel, N. L. Bray, S. Puente, P. Melsted, and L. Pachter. Differential analysis of RNA-Seq incorporating quantification uncertainty. Nat. Methods, 14:687–690, 2017.

[56] M. I. Love, S. Anders, and W. Huber. Analyzing RNA-seq Data with DESeq2, 2017. R Package Reference Manual.

[57] B. Li and C. N. Dewey. RSEM: Accurate transcript quantification from RNA-Seq data with or without a reference genome. BMC Bioinformatics, 12:323, 2011.

[58] D. Szklarczyk, K. Nastou, M. Koutrouli, R. Kirsch, F. Mehryary, R. Hachilif, D. Hu, M. E. Peluso, Q. Huang, T. Fang, et al. The STRING database in 2025: Protein networks with directionality of regulation. Nucleic Acids Res., 53:D730–D737, 2025.

[59] S. X. Ge, D. Jung, and R. Yao. ShinyGO: A graphical gene-set enrichment tool for animals and plants. Bioinformatics, 36:2628–2629, 2020.

[60] N. A. O’Leary, M. W. Wright, J. R. Brister, S. Ciufo, D. Haddad, R. McVeigh, B. Rajput, B. Robbertse, B. Smith-White, D. Ako-Adjei, et al. Reference sequence (RefSeq) database at NCBI: Current status, taxonomic expansion, and functional annotation. Nucleic Acids Res., 44:D733–D745, 2016.

[61] E. T. Wang, R. Sandberg, S. Luo, I. Khrebtukova, L. Zhang, C. Mayr, S. F. Kingsmore, G. P. Schroth, and C. B. Burge. Alternative isoform regulation in human tissue transcriptomes. Nature, 456:470–476, 2008.

